# Quantifying defective and wild-type viruses from high-throughput RNA sequencing

**DOI:** 10.1101/2024.07.23.604773

**Authors:** Juan C. Muñoz-Sánchez, María J. Olmo-Uceda, José-Ángel Oteo, Santiago F. Elena

## Abstract

Defective viral genomes (DVGs) are variants of the wild-type (wt) virus that lack the ability to complete an infectious cycle independently. However, in the presence of their parental (helper) wt virus, DVGs can interfere with the replication, encapsidation, and spread of functional genomes, acting as a significant selective force in viral evolution. DVGs also affect the host’s immune responses and are linked to chronic infections and milder symptoms. Thus, identifying and characterizing DVGs is crucial for understanding infection prognosis. Quantifying DVGs is challenging due to their inability to sustain themselves, which makes it difficult to distinguish them from the helper virus, especially using high-throughput RNA sequencing (RNA-seq). Accurate quantification is essential for understanding their interactions with their helper virus. We present a method to simultaneously estimate the abundances of DVGs and wt genomes within a sample by identifying genomic regions with significant deviations from the expected sequencing depth. Our approach involves reconstructing the depth profile through a linear system of equations, which provides an estimate of the number of wt and DVG genomes of each type. Until now, *in silico* methods have only estimated the DVG-to-wt ratio for localized genomic regions. This is the first method that simultaneously estimates the proportions of wt and DVGs across RNA sequencing of the whole genome.

**Availability and implementation:** The Matlab code and the synthetic datasets are freely available at https://github.com/jmusan/wtDVGquantific.

## 1 Introduction

During viral replication, a plethora of varying copies of the genome are produced, including point mutations and hypermutations, insertions, deletions, and genome recombinations [1]. Defective viral genomes (DVGs) are a subset of them characterized by requiring the presence of the wild-type (wt), or helper, virus to complete a viral infectious cycle. Their production has been traditionally associated with random errors in the viral replication but recent studies suggest their generation is not entirely random but mediated by viral and host factors [2–4]. Some authors have even proposed that they may provide a fitness advantage to the virus [5]. Since first described in influenza A virus [6], examples of DVG-producing virus has been documented in positive and negative single- and double-stranded RNA viruses and retroviruses [1, 4]. DVGs influence the course of infection in several ways: (*i*) by interfering with wt replication as they can compete for cellular resources during replication; (*ii*) by modulating the immune response of the host as some DVGs might express viral proteins that can interact with the host’s immune system affecting the severity and duration of the infection; and (*iii*) by imposing evolutionary pressures as they can recombine with functional viral genomes, leading to the generation of novel viral variants with altered properties, including enhanced or reduced virulence [1].

Because of these interfering and immune-stimulation characteristics, the potential of DVGs in therapeutic applications has been explored. The so-called therapeutic interfering particles (TIP) have been shown to successfully reduce the accumulation and transmission of flavivirus [7], Dengue virus [8] and SARS-CoV-2 [9–11] in cell cultures and model animals. The number of mathematical models trying to disentangle the dynamics between the wt and its DVGs is also increasing [12, 13]. Quantifying both wt virus and DVGs is important for studying infection progression but becomes a computationally challenging problem when we do not focus on a particular DVG but rather the interest is to characterize genome-wide generated DVGs coexisting in a viral population. Most viruses are readily quantified by simple plaque assay in susceptible cells. However, since DVGs do not complete a full infection cycle, their quantification by plaque assays is not feasible and more laborious biochemical techniques have traditionally been used for their isolation and quantification. High-throughput sequencing (HTS) techniques, along with new bioinformatic pipelines applied to HTS RNA sequencing (RNA-seq) samples, had opened the possibility of identifying the DVG component of the mutant swarm. Despite the revolution offered by short-read RNA-seq, it also imposes some errors and biases. Improved strategies such as direct RNA Nanopore sequencing [3] and circular sequencing [14] have improved the detection of DVGs. However, short-read RNA-seq still remains the most commonly employed technique.

Briefly, a short-read RNA-seq process consist in the extraction of the total RNA from a sample, followed by their fragmentation in short fragments, the preparation of the library (the fragments are fused to known small sequences) and its sequencing by amplification with next-generation sequencing (NGS) methods. The resulting sequencing reads are saved into fastq format files. If a reference genome is available for the virus of interest (as is usually the case), the reads are mapped to the reference and an alignment file (in SAM and BAM formats) is generated [15]. This alignment file can be interrogated to know how many reads align at each position of the genome (herewith obtaining the depth profile), being this a proportional measure of the abundance of each part of the genome in the original sample. Software such as ViReMa-a [16], DI-tector [17], DVG-profiler [18], or VODKA2 [19] can help to identify the abnormal junctions that characterize DVGs in the short reads of the RNA-seq data. All this algorithms align the short reads sequenced against the wt genome and identify the recombination events by marking the dis-joining points, *i.e*., the two positions that are non-contiguous in the wt but that are adjacent in the DVG. This two positions are commonly dubbed as the breakpoint (BP) and the re-initiation site (RI). We developed DVGfinder [20], a meta-search tool that integrates the two most widely used search algorithms, ViReMa-a and DI-tector, unifying their outputs and adding information to characterize the complexity of the DVG population. DVGfinder provides estimates on the diversity and abundance of DVGs based on the number of reads mapping the recombination event.

So far, a measure of local abundance has been used to estimate the DVG abundance in the sample, being the quotient between reads mapping the chimera and total mapped reads the most employed one [2,3,7,21,22]. However, the estimated number of complete genomes for each predicted DVG remains unexplored. In this study, we propose a method to obtain the ratio between wt and DVGs in an RNA-seq sample using as starting point the output from DVGfinder, the depth information by position and some linear algebra. We believe that this ratio, complemented with results from plaque forming assays (to estimate wt virus), provide approximate values for the number of DVGs in the samples.

In Section 2 the method will be introduced as well as the inputs needed for its application. In Section 3 a step-by-step synthetic example will be generated and analyzed to evaluate the discrepancies obtained in the quantification of the DVGs. Finally, Section 4 presents a case study using data from Hillung *et al*. [23].

## 2 Method

### 2.1 Statement of the problem

The serial dilutions method used to estimate the virus concentration in a sample is unable to provide information about the DVG content we are interested in. We propose a method to determine the number of DVGs from RNA-seq data by including the information present in the alignment depth profile. We begin by carrying out an exploratory spectral analysis of the depth profile which leads us to conclude that there are no particular enhanced spectral modes that could eventually be associated to DVG structures. Next, we take advantage of some results provided by DVGfinder that can be readily interpreted as DVGs of elementary character. We denote them as eDVGs and will be the instrumental basis of the method to assess the amount of DVGs present in a sample. The idea is to consider the experimental depth profile of the sample as a weighted aggregate of a number of these elementary DVGs. The weights account for the relevance of specific DVGs in the sample. This procedure leads us eventually to establish an empirical mode decomposition of the depth profile function that estimates the spectrum of eDVGs of each type present as a proxy of the actual DVGs and wt content.

### 2.2 The Walsh spectrum of the depth profile

DVGfinder uses as starting point the fastq files from the HTS to obtain the diversity of DVGs in the sample and, as a by-product of the procedure, a per-site depth profile 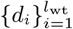, being *i* the base number, *d*_*i*_ the depth value at that base and *l*_wt_ the length of the wt virus sequence. In order to ascertain whether some spectral modes are particularly enhanced in the depth profile, we look into its spectral content. The idea is to consider the values *{d*_*i*_*}* as a discretely sampled function and using a spectral method to analyse it. We have discarded the option of Fourier analysis because this case is greatly affected by the Gibbs phenomenon resulting from the piece-wise constant nature of the profile. In this situation it is more convenient to expand the depth function into Walsh series [24] because the Walsh modes resemble square waves on the unit interval. The functions *W* (*k, x*) of the Walsh basis are non-periodic and take amplitude values *±*1. The index *k* stands for the number of zero-crossings and defines the quasi-oscillation mode. Walsh functions with *k* even (odd) are symmetrical (anti-symmetrical) with respect to *x* = 1*/*2. The fundamental mode is simply *W* (0, *x*) = 1.

To associate the measured *d*_*k*_ values of the depth profile with a real function *d*(*x*) defined in the unit interval we map the index *k* into the real number *k/l*_wt_ ∈ [0, 1], which will play the role of *x*, and define the discretely sampled function *d*(*k/l*_wt_) = *d*_*k*_, in the unit interval.

The functions *W* (*k, x*) form a normal basis with respect to the scalar product

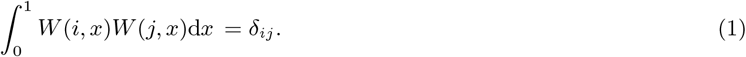

The next step consists in expanding *d*(*x*) in terms of the functions of the Walsh basis. The Walsh expansion up to order 2^*n*^, with *n* an integer, reads formally

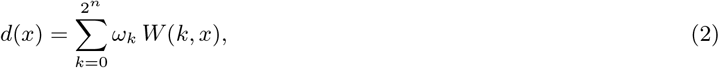

with

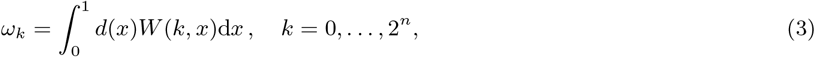

the Walsh amplitudes.

Two instances of Walsh spectra for the absolute amplitudes |*ω*_*k*_|, (3), are given in Fig. 1. They correspond to two instances made from synthetic data (upper panel) and from real data (bottom panel), whose details will be provided in Sections 3 and 4, respectively.

**Figure 1:**
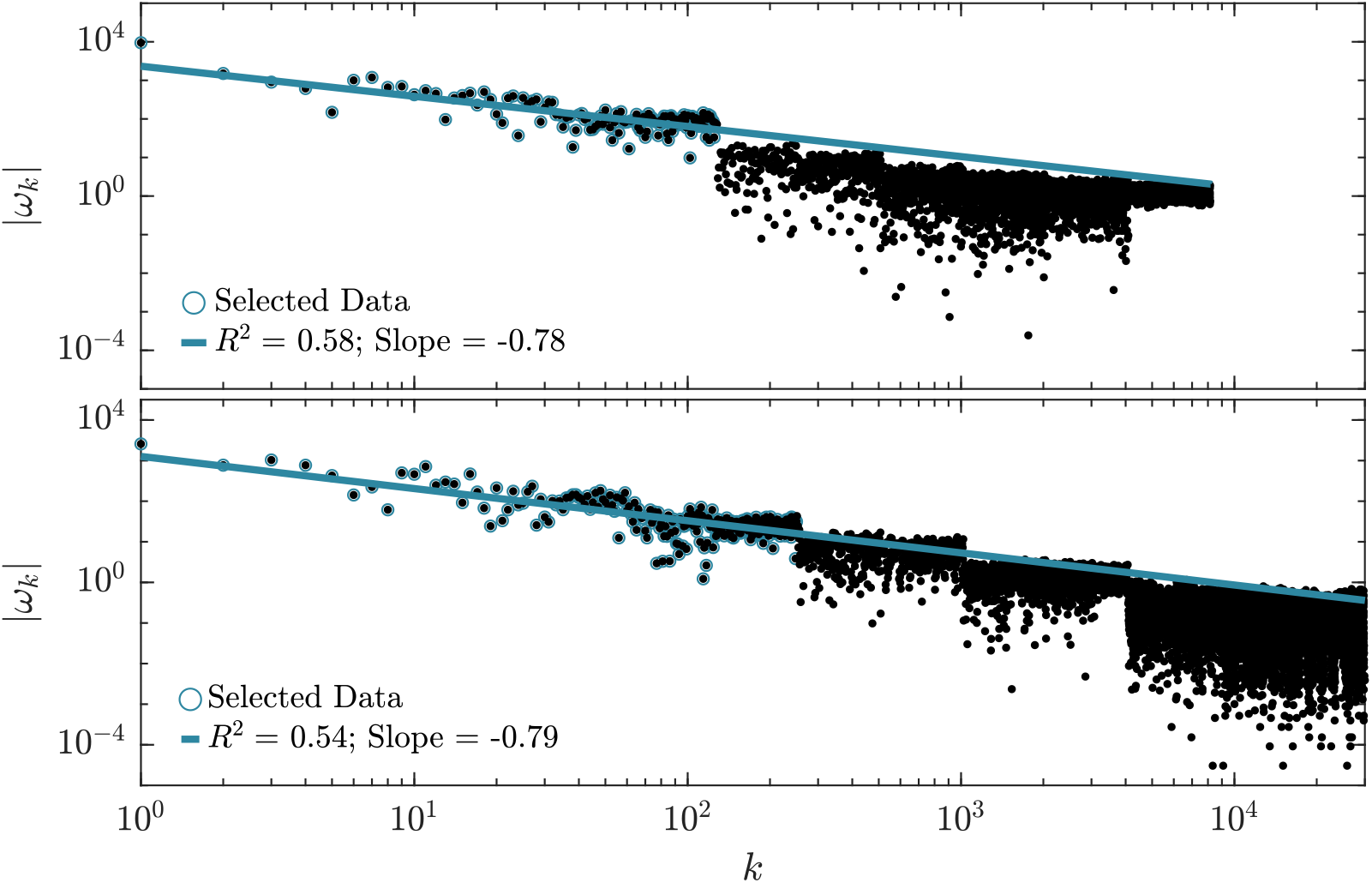
Walsh spectrum for the synthetic (top panel) and real (bottom panel) examples in Sections 3 and 4. Blue circles correspond to the data that have been used for the linear fit (blue lines).

For the time being, it suffices to observe the approximate power law character of the distributions of Walsh weights |*ω*_*k*_| with respect to the zero-crossing index *k*. This outcome points out that no Walsh zero-crossing mode, or combination of them, is enhanced in the spectra (no peak). In the case of Fourier spectra, power law distributions are associated to noise. More specifically, to white noise whenever the slope is minus one and to colored noise otherwise. Since the zero-crossing mode index *k* plays in Walsh spectra the role that the frequency plays in Fourier spectra, we are then lead to argue that the profiles in Fig. 1 correspond (approximately) to colored noise. Thus, the conclusion of the exploratory analysis is negative: no elementary combination of Walsh modes seems to characterize the presence of DVGs or, at least, cannot be detected by this technique. We will return to this issue below. Next, we describe a successful alternative to analyze the content of DVGs in the depth profile.

### 2.3 The elementary DVGs

The estimation of the DVGs content in a sample that we propose relies on the introduction of a set of elementary DVGs depth profiles. Elementary refers to the very basic shape of their depth profiles, namely made of a few segments of piece-wise constant functions, as illustrated in Fig. 2, that we describe next. Whereas the depth profile of the wt genome is represented by a constant unit function (Fig. 2), the eDVGs profiles are not constant because they describe one of these three local molecular processes: deletion, insertion or copyback (cb), that take place in one, and only one, region of the genome and whose BP and RI sites are provided by the DVGfinder diversity table (*e.g*., Table D1). The eventual eDVGs resulting from the simultaneous action of several recombination events are not contemplated in the present analysis. The depth profiles we are dealing with, associated to each eDVG, are also illustrated in Fig. 2. As already mentioned above, the wt depth profile is constant with value 1 because the wt virus contributes to the depth profile with one count per base. Deletions make the profile to have one, and only one, arbitrary length sequence filled with zeroes, namely zero counts per base. For insertions, the variable length sequence is filled with 2s, namely two counts per base. The components of cb DVGs are the juxtaposition of three sequences of variable length filled with values 0, 1 and 2, respectively (Fig. 2).

**Figure 2:**
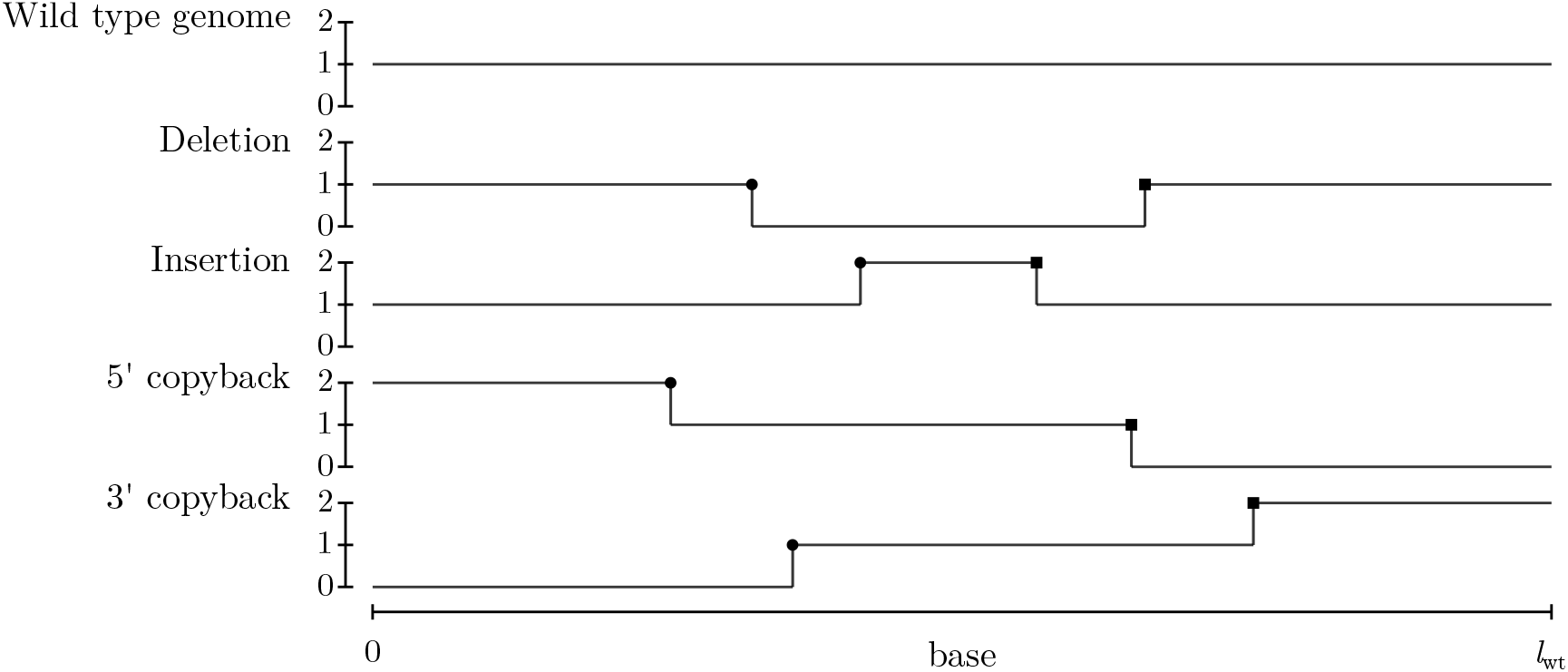
An instance of elementary depth profiles: wt, deletions, insertions and copybacks (cb). The dis-joint points are labeled as “start” (circles) or “end” (squares) depending on their position with respect to the wt reference positive genome, as usually done for non-stranded RNA-seq analysis. Deletions (insertions) are characterized by a lack (duplication) of part of their genome resulting in a the depth profile with null (double) coverage between its dis-joint points. Copybacks are characterized by a loop secondary structure between their dis-joint positions in which their contribution will be the same that the wt virus in that region. The 5’cb (3’cb) will double contribute to the coverage between the 5’ (3’) site and the “start” (“end”) position and will not contribute to the coverage between “end” (“start”) and 3’ (5’) positions. In practice, these are discrete functions sampled at l_wt_ base locations.

The horizontal profile character of a wt is broken by the presence of DVGs in an experimental sample. Our main assumption is then that the depth profile of the sample can be comprehensibly described in terms of the set of eDVGs provided by DVGfinder altogether with the very wt profile. The way this can be achieved is described next.

### 2.4 The determination of the eDVG spectrum

Given the data of a virus sample provided by DVGfinder, *i.e*., the set of eDVGs and the depth profile of the sample; the problem that we solve is the determination of the weight that each eDVG has in that sample. We refer to this set of weights as the eDVG spectrum of the sample. We are then able to provide not only an estimate of the usual DVG-to-wt ratio of the sample, but also a non-local measure of the relevance of the wt virus and each eDVG, which is an alternative that goes beyond the common local *reads*.

To determine the eDVG spectrum we interpret the *l*_wt_ values that define the depth profile of the sample, the wt and each eDVG as the coordinates of vectors in a phase space of dimension *l*_wt_. This is a geometric representation of all the experimental information. Every axis of the phase space corresponds to a base of the reference genome. In turn, each point represented in phase space stands for a whole depth profile of: (*i*) the sequenced sample, say vector 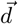, (*ii*) the wt, whose vector 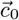 has all the coordinates equal to one, or (*iii*) the elementary DVGs, with vectors 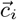, and *i* = 1, …, *D*. All these vectors are located in the first orthant of the phase space because their components are non-negative numbers. We pose then the following task: to determine the linear combination (in the sense of Algebra), with non-negative coefficients, of the *D* + 1 vectors 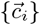 that best mimics the virus vector 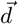, namely

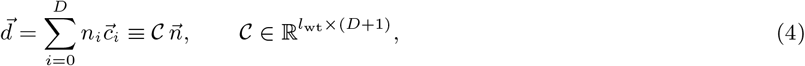

where we have defined the vector of coefficients 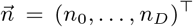. We will refer to 𝒞 as the contribution matrix. Interpreted as a linear algebraic system of dimension *l*_wt_ with *D* + 1 unknowns *n*_*i*_, (4) is an over-determined linear system, *l*_wt_ ≫ *D*. Because the number of equations exceeds the number of unknowns, we can only get a determination in the sense of Least Squares (LS), with the constraint that the values of *n*_*i*_ have to be non-negative. We have used the built-in Matlab routine *lsqnonlin* to this end. The ratio 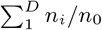 is the estimated proportion DVG-to-wt in the sample and the set *{n*_*i*_*}* is the eDVG spectrum.

Alternatively, to solve equation (4) one can expand the depth profile, *d*, the wt, *c*_0_, and the eDVGs, *c*_*i*_, into the Walsh basis and use then the scalar product given by equation (1) to determine the coefficients *c*_*i*_. We have checked that this procedure leads to similar spectra, albeit in a slightly more indirect way. It is noteworthy that the Walsh spectra of the eDVGs themselves do not exhibit the power-law character of the depth profile spectra shown by the synthetic and the real samples in Fig. 2.

## 3 Synthetic example

A comprehensive example of the method is presented using a synthetic dataset. Section 3.1 describes the way the dataset has been generated. Next, in Section 3.2, a procedure to noise reduction and artifact removal is presented, which is of outstanding interest when analyzing real datasets. Eventually, the processed dataset is used to get the eDVG spectrum.

### 3.1 Synthetic dataset

We have used SDgenerator, a free software developed by our group (available to download from https://github.com/MJmaolu/SDgenerator/tree/main), to generate the synthetic dataset (see Appendix A for details) according to the parameters in the first four columns of Table 1. Six defectives genomes with nominal proportions 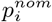 have been introduced. In addition, we have added some noise to simulate intrinsic biological and processing noise (alignment and DVG detection errors).

**Table 1:** SDgenerator input table and estimated results. The four leftmost columns of the table define the genomes present in the synthetic dataset where 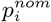 stands for the nominal proportion of each genome. The two rightmost columns correspond to the outcome of the quantification process: p_*i*_ stands for the eDVG estimated proportion and Δ_*i*_ is the relative discrepancy.

| Specie | BP | RI | $p_i^{\text{nom}}$ | $p_i$ | $\Delta_i$ (%) |
| --- | --- | --- | --- | --- | --- |
| wt | - | - | 0.30 | 0.2953 | 1.5 |
| Deletion | 2000 | 3000 | 0.10 | 0.1008 | 0.8 |
| Deletion | 7000 | 9000 | 0.15 | 0.1515 | 1.0 |
| 3' cb/sb | 8000 | 8750 | 0.05 | 0.0498 | 0.4 |
| 5' cb/sb | 3100 | 2200 | 0.05 | 0.0517 | 3.4 |
| Insertion | 3500 | 5500 | 0.10 | 0.1047 | 4.7 |
| Insertion | 4100 | 4900 | 0.25 | 0.2462 | 1.5 |

The DVGfinder outcome for the depth profile of the synthetic dataset is shown in Fig. 3. Each vertical line represents a recombination event (BP or RI positions) of the DVGs listed in Table 1. Ideally, the points in between should have the same height, giving rise to plateaus that, in practice, are blurred by noise and artifacts. Note also the side effect at the beginning and the end of the genome, as well as transition profiles. Next, we explain a procedure that allows to clean up the depth profile prior to the LS analysis.

**Figure 3:**
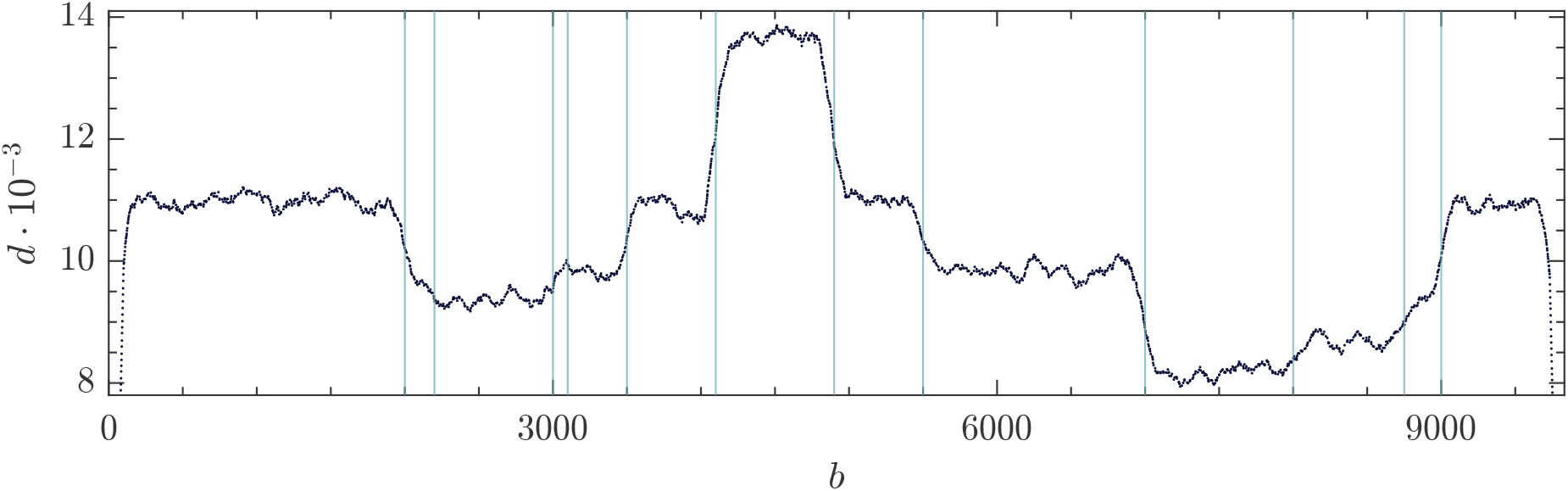
Depth profile of the synthetic dataset. Vertical lines indicate the presence of a disjoint or recombination site.

### 3.2 Artifact removal and noise reduction in the depth profile

Ideally, points between two consecutive vertical lines in Fig. 3 should be plateaus, so that have the same coverage height. Thus, there is a need to clean up the data. The goal is to give a determination of the height of each plateau between vertical lines in Fig. 3. To this end, we define variability thresholds in each interval that rule out extreme values in the profile. We determine the median 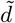 in each plateau-like interval using only its central 80% points in order to avoid border effects and minimize the effect of outlier values. After that, all the data *d*_*i*_ of each interval are mapped to the relative depth deviation with respect to the plateau’s median

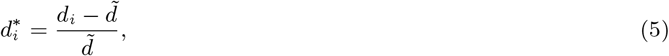

where *i* stands for the base in that interval. The relative variations for every base are represented in Fig. 4. The 95% quantile of the distribution of the relative distances is then used to set the right cutoff 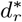. The left cutoff 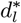 is then symmetrically fixed with respect to the median of the distribution 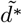, namely, 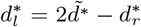. The horizontal red lines in Fig. 4 locate the cutoffs. Data outside these limits are therefore discarded for the constrained LS analysis.

**Figure 4:**
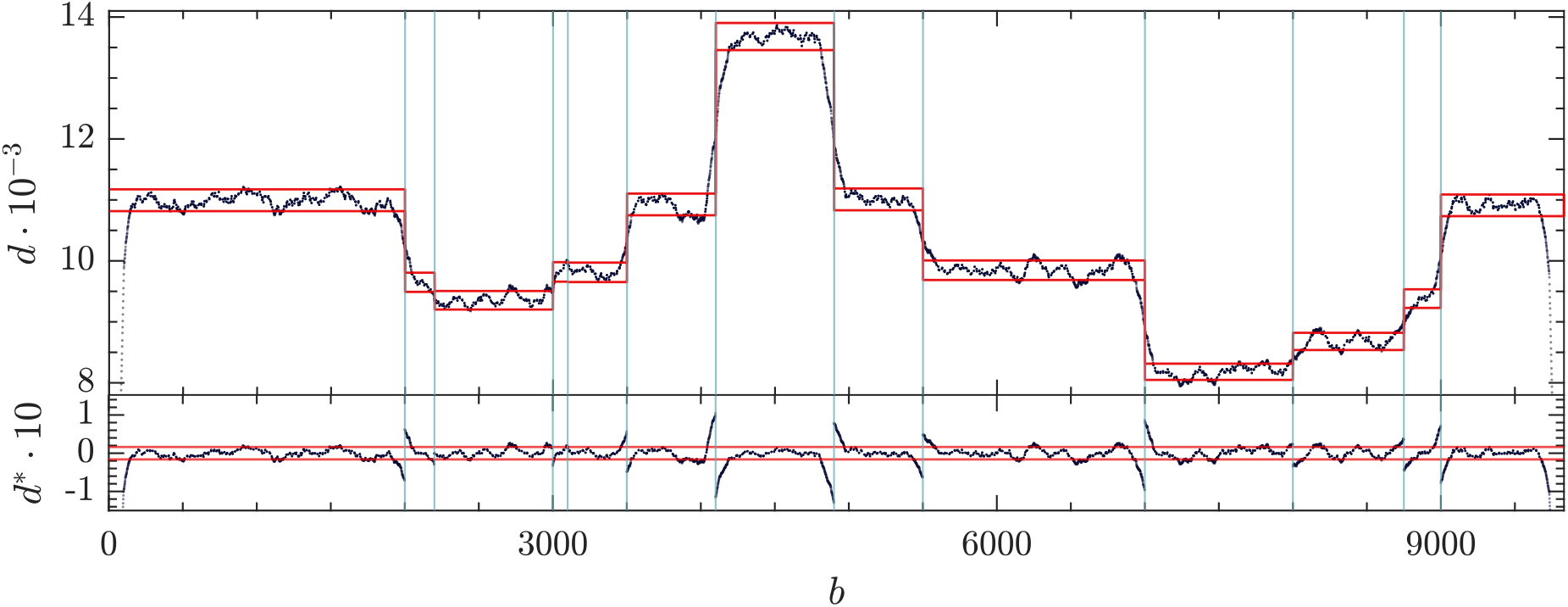
(Top) Depth profile of the synthetic dataset (black dots). Vertical lines locate DVG limits (BP and RI) as DVGfinder output. Horizontal red lines bound the allowed points used to the analysis. (Bottom) Relative depth deviation 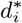 (5). Horizontal red lines locate the cutoffs.

### 3.3 eDVG spectrum estimate

Once the depth profile in the dataset has been pre-processed, the next step is to solve the linear system in equation (4) whose effective dimension 𝓁 has decreased, 𝓁 *< l*_wt_, because of the points dropped out in the datafile debugging described in the previous Section. The contribution matrix, *𝒞*, for this example can be found in Appendix B. We have used the built-in Matlab routine *lsqnonlin* that carries out the LS fit constrained to non-negative solutions, *n*_*i*_ ≥ 0, *i* = 0, …, *D*. The outcomes of the analysis are shown in Table 1, where 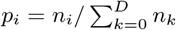, with *i* = 0, …, *D*, stands for the proportion of wt virus (*p*_0_) and eDVGs (*p*_*i*_, *i >* 0) in the sample. The last column in Table 1 shows the relative error with respect to the nominal value, 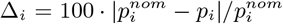. The outcomes *{n*_*i*_*}* are real non-negative numbers. In practice, we deal with the number of genomes of each species which is estimated as the nearest integer. The backward reconstruction of the depth profile from the LS estimates *{n*_*i*_*}* in Fig. 5 (green horizontal line) shows that the piece-wise constant function based on the six eDVGs plus the wt provides good agreement with the experimental depth profile.

**Figure 5:**
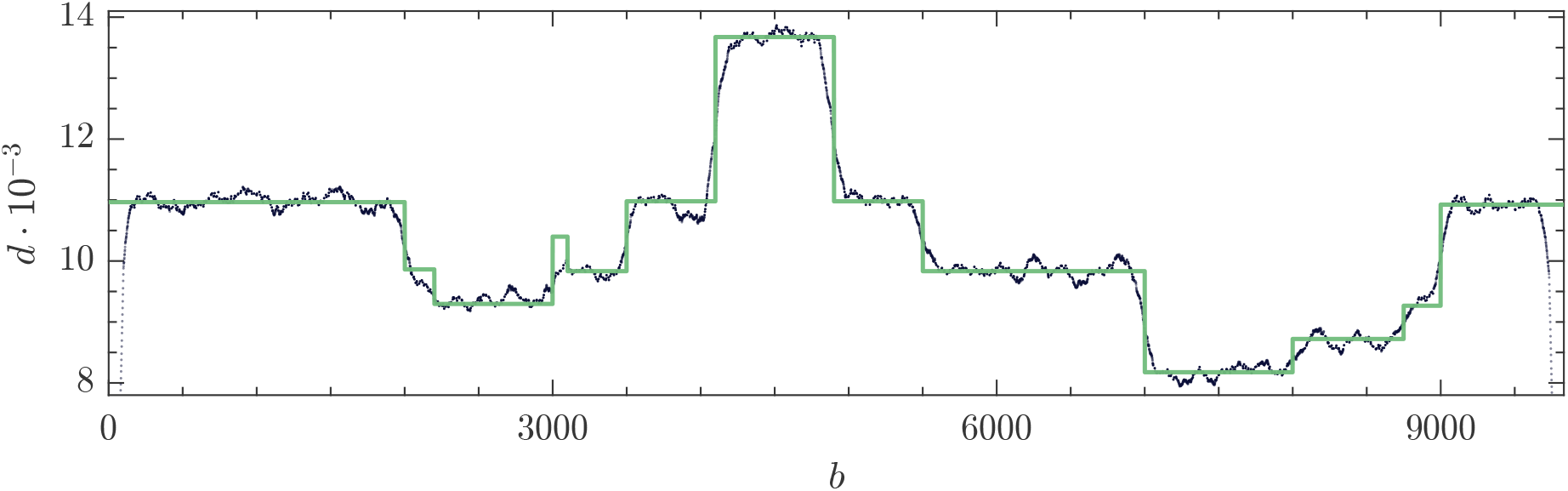
Synthetic data depth profile (black dots) and the estimated depth profile (solid green line).

## 4 Case study: the eDVG spectrum of HCoV-OC43 in a cell culture

To elucidate the dynamics of DVG accumulation in evolving betacoronaviruses, Hillung *et al*. [23] performed serial passages of HCoV-OC43 populations in baby hamster kidney cells (BHK-21). Inoculum sizes varied between passages within two broad but disjoint intervals that can be roughly defined as low and high multiplicities of infection (MOIs). At every passage, viral particles were estimated using as a proxy the number of plaque forming units (PFUs). Some passages were sequenced by HTS RNA-seq. Here, to illustrate the application of our DVG quantification method, we will focus in the RNA-seq data obtained from the first passage.

Depending on the software employed to obtain the set of DVGs in the sample and the depth profile, it may be necessary to pre-process the data to retain only those DVGs that significantly contribute to the depth profile. Appendix C presents the algorithm used in instances where the depth profile and the set of DVGs was generated using DVGfinder.

The post-processing of the raw data is summarized in Fig. 6. The depth profile is represented as black dots in the top panel. Noise and artifacts are apparent. The red horizontal lines determine the bands whose data are used in the constrained LS solution of equation (4). The vertical blue lines indicate the BP and RI positions of the remaining eDVGs after the pre-processing steps. The purple vertical lines indicate the subgenomic positions of the wt virus. The relative depth deviation *d*^*^ (5) is represented in the bottom panel.

**Figure 6:**
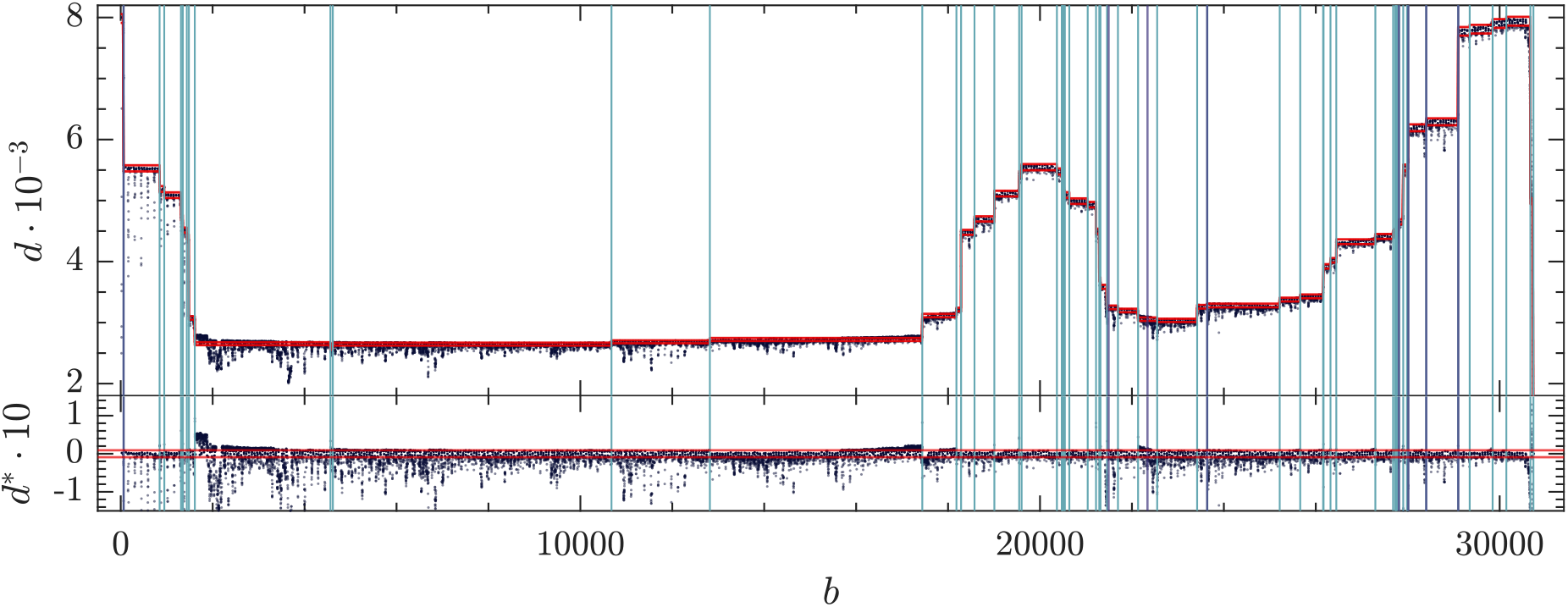
(Top) Depth profile of the real sample with the noise limits computed at each plateau (red lines). Vertical blue lines indicate the BP and RI positions of the remaining eDVGs after the pre-processing steps. Purple vertical lines indicate the subgenomic positions. (Bottom) Relatives distances d^*^ and cutoffs (red lines).

The outcome of the constrained LS analysis is reported in Table D2 and in Fig. 7. The piece-wise constant line is the reconstructed depth profile, to be compared with the raw depth profile (black dots). According to these results, 28.8% of the sample corresponds to wt virus (including the subgenomics), 70.2% to deletions, 1.0% to insertions and 0.0% to cb. Therefore, in this sample of HCoV-OC43 the ratio DVG-to-wt is 2.47.

**Figure 7:**
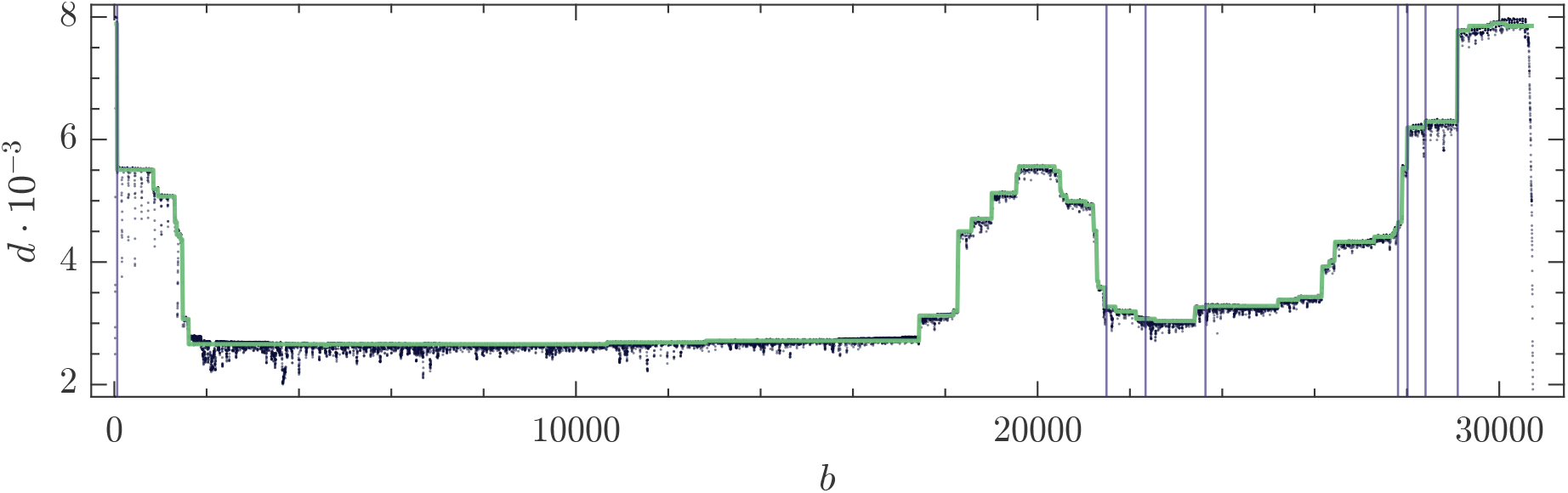
Real sample depth profile (black dots) and the estimated depth profile (solid green line). Purple vertical lines stand for the subgenomic RNAs positions.

The number of genomes *n*_*i*_ is estimated as the nearest integer value to the constrained LS solution. A comparison between the local measure given by *reads* and the estimates of the method is in Fig. 8. In order to allow a faithful comparison of both quantities, they have been normalized to sum one.

**Figure 8:**
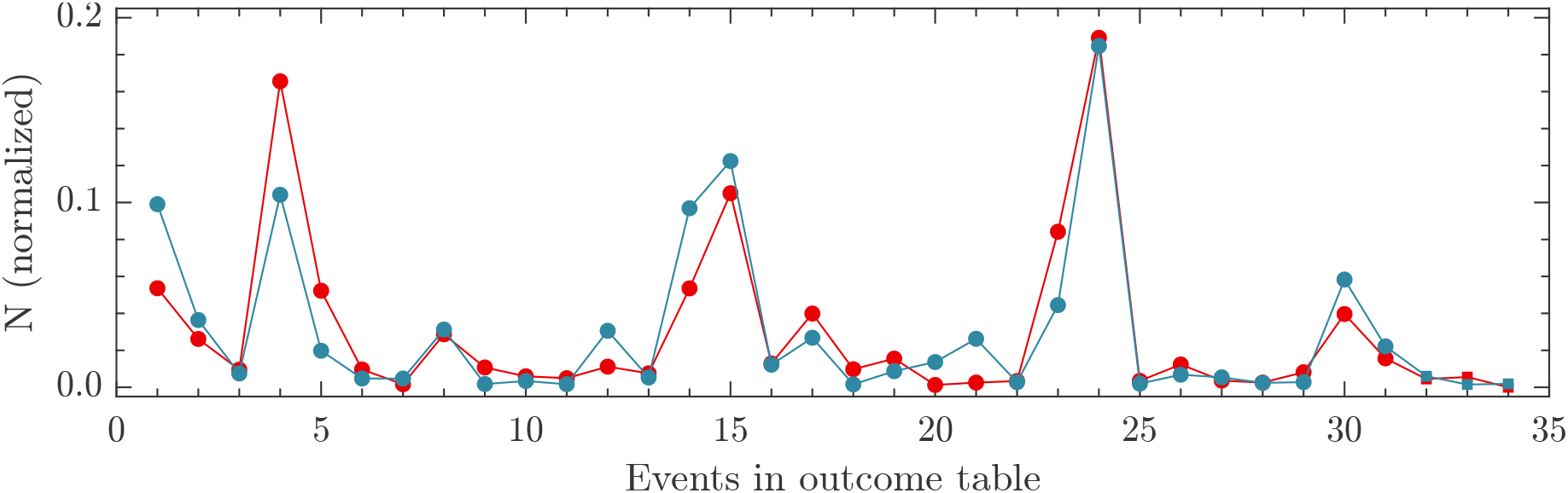
Outcomes of the analysis of the real sample. Number of eDVGs (red) and ViReMa-a reads (blue) normalized to sum one. Dots stand for deletions and squares (rightmost part) for insertions. No copybacks detected in this sample.

## 5 Discussion

Any theoretical result is prisoner of the hypotheses and approaches formulated to achieve it, and this work is not an exception. Key assumptions essential for achieving the presented results include the single-event eDVG hypothesis and the unequivocal presence of DVGs in the sequencing depth profile. In the test with synthetic dataset, in which both assumptions were satisfied simultaneously, the algorithm worked successfully. When applied to real data, the depth profile was correctly reconstructed and the estimated genomes correlated well with the ViReMa-a reads obtained using DVGfinder (linear correlation with slope 0.86 *±* 0.07 with *R*^2^ = 0.81). The consideration of eDVGs with multiple events conveys adding new rows to the contribution matrix and should be taken into account whenever the depth profile cannot be properly recovered. Some cautions should be taken when insertions or deletions with |BP−RI| small appear (typically a few bases), as their depth profiles are extremely similar to wt virus’. Adding a filter in which only events bigger than certain number of bases are considered avoids this problem at the price of adding some genome counts to the wt virus that do not actually belong to it.

The usefulness of the method proposed here relies in the assumption that the sequencing depth at each nucleotide position represents the proportions of the original sample genomes. We have optimized the method for data sequenced from total RNA extractions and amplified with random hexamers. The sources of bias this type of data can present, from sample preparation to the aligner used have been discussed in depth [25–27]. Regarding the pre-processing steps and the noise reduction, more sophisticated strategies could be implemented. In the present case we opted to keep the simplest approach to avoid distracting the reader from the main topic. However, if the noise sources are known (better knowing the samples’ and the aligner’s noise) some statistical distribution of the noise could be derived and used to better estimate which data points are outliers or more affected by the noise.

This work forms part of a broader project aimed at studying the dynamics of appearance and accumulation of DVGs and their interaction with wt helper virus and the host. A series of interconnected experiments were conducted to tackle different aspects of this dynamical process. Chronologically, initial experimental results revealed variations in viral load estimated through plaque assays among passages hinting at the possibility of periodic interaction with DVGs. Simultaneously, a theoretical model was developed [12], hypothesizing that these oscillations were not due to the presence of DVGs but rather experimental noise. According to the model, the wt virus and DVGs reached a certain equilibrium during serial passages at different MOIs, jointly varying their post-infection values. Additionally, stability analysis of the model yielded quasineutral planes and sensitivity to initial conditions, potentially explaining observed variations in [23]. This simple theoretical model was validated by fitting experimental data obtained from another study [2], motivating us to refine the estimation of defective particles throughout passages and compare them with estimated PFUs to shed some light on these observed oscillations. The present algorithm was developed to simultaneously determine the number of genomes associated with DVGs and wt virus. Assuming that the plaque assay provides a reliable estimate for infectious virions, *V*, proportional to wt genomes within cells, it was postulated that virions encapsidating defective genetic material, *V*_*D*_, would have the same proportionality factor. Thus, the number of DVGs could be estimated using the ratio *V*_*D*_ = (DVG*/wt*) *· V*, where *V* is associated to PFUs. Applying this algorithm to all samples from the HCoV-OC43 infection experiment in BHK-21 at low and high MOI, an estimate of viral particles encapsulating DVGs was obtained. Notably, experimental validation of this process is extremely challenging as DVGs do not complete a full infection cycle, preventing plaque assays. Fig. 1b in [12] presents estimates for *V* through plaque assays alongside estimates for *V*_*D*_ by the present method. Oscillations characteristic of a Lotka-Volterra host-parasite arms race were not observed, consistent with results found in other associated studies [2, 12, 23].

In summary, we have proposed a method to estimate the proportion of DVGs and wt within a sample sequenced by RNA-seq short-reads, prior amplification of total RNA with random hexamers. Amplicon-based amplification techniques, which involve the targeted amplification of specific regions of the viral genome, are not recommended for characterizing DVGs due to the risk of losing crucial recombination events. On the other hand, Cir-seq and long-reads sequencing techniques are being implemented in the field with increasing success and some parts of the method could be useful adapted to their particularities, although it is beyond the scope of this work. We used the results from DVGfinder but any DVG detection tool can be employed as far as it provides DVG types and the recombination points. In any case, a depth by position file is also needed. In case DVGfinder is run, we recommend using the intermediate file “*{*sample*}* sorted mapped depth.txt”. Subgenomic RNAs, a canonical type of deleted genomes through some virus translate their structural proteins, can be also be quantified by this method. We expect the method presented to be useful for most of the data already produced.

## Author contributions

The authors confirm their contribution to the paper as follows: study conception and design: J.C.M.S., M.J.O.U., J.A.O.; analysis and interpretation: J.C.M.S., M.J.O.U., J.A.O., S.F.E.; writing—original draft preparation: J.C.M.S.; writing—review and editing: J.C.M.S., M.J.O.U., J.A.O., S.F.E.; All authors have read and agreed to the published version of the manuscript.

## Acknowledgements

Many computations were performed on the HPC cluster Garnatxa at I^2^SysBio (CSIC-UV). J.C.M.S. and S.F.E. acknowledge the support of the Santa Fe Institute, where part of this research was developed.

## Appendix

### A SDgenerator

SDgenerator v.2 is a Python program developed to simulate short-read RNA-seq samples formed by controlled populations of wt virus and its associated DVGs. The algorithm takes as input the reference genome (*i.e*., the wt virus) and a table with the DVG composition with the next information: DVG type, BP, RI, and proportion of each DVG in the sample. The user can control for the library size (*N*_*T*_, total reads) and the length of the reads (*L*) of the synthetic sample. The code is available at *https://github.com/MJmaolu/SDgenerator/tree/main*.

**Figure A1:**
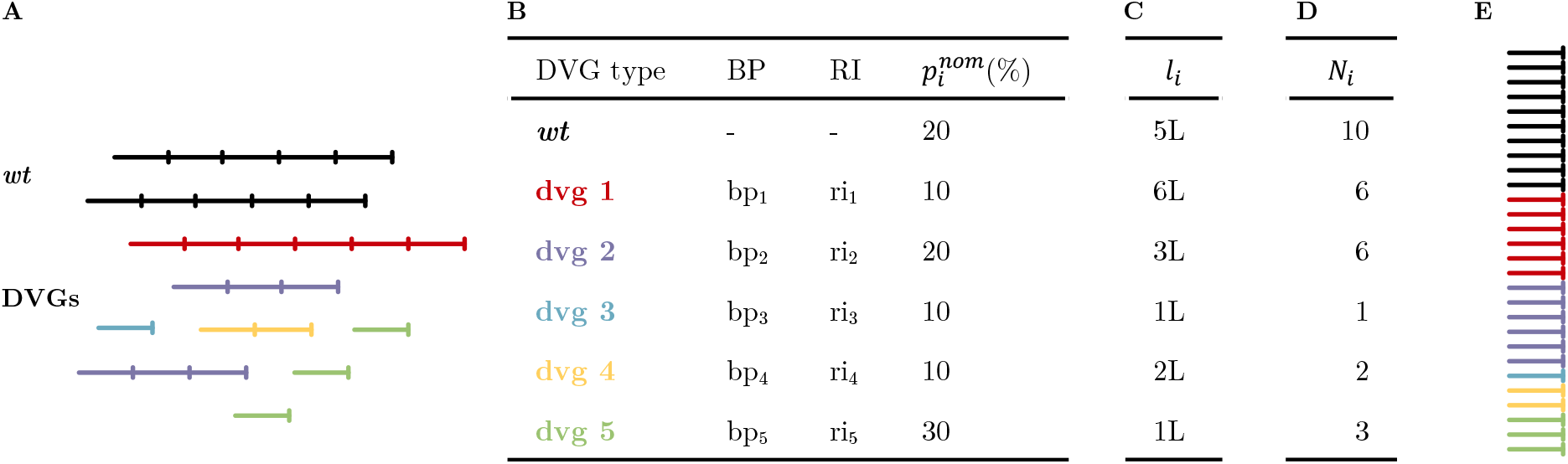
Schematic pipeline of SDgenerator. (A) Expected genome composition. For simplicity a population of only six different genomes is represented. Black genomes are the wt and colored genomes represent different DVGs. The segments indicate arbitrary unites of length. (B) Genomes in (A) represented as input table for SDgenerator, the number of genomes of each type are indicated as proportion in the total sample 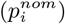. (C) The reconstructed length of each genome (l_*i*_) is calculated as in [20]. (D) Finally, the number of reads to generate a sample with N_*T*_ = 28 and L = 1 is calculated A1 and (E) the program launches the generation of synthetic reads with wgsim and concatenates the results in a single fastq file.

Being *N*_*T*_ the desired total number of reads with length *L, N*_*T*_ the total number of genomes, *l*_*i*_ the length of each genome with proportion 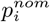, the total number of reads associated with each genome *N*_*i*_ is:

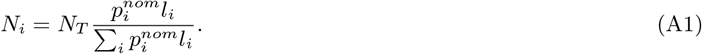

*Proof* :

User inputs: *N*_*T*_ and 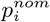 (from which *l*_*i*_ is derived knowing BP and RI). The number of reads to assign to each genome *g*_*i*_ (*N*_*i*_) is the amount of copies of that genome (*n*_*i*_) times its length (*l*_*i*_) divided by the length of the read (*L*):

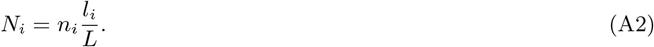

The number of genomes of each specie (*n*_*i*_) can be obtained knowing the total number of genomes and the proportion of the sample corresponding a *g*_*i*_:

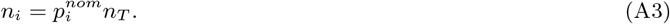

The total number of genomes in the sample can be obtained starting from the total number of reads (*N*_*T*_) and expressing it as the sum of the reads assigned to each individual genome. If this individual contribution is written in terms of the proportion of each genome in the sample and the total number of samples then *n*_*T*_ can be isolated as follows:

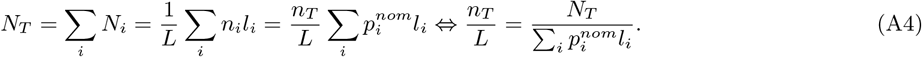

From (A2) -(A4) one can easily obtain (A1).

Once each genome has been reconstructed (an saved in fasta format) and its *N*_*i*_ has been calculated, SDgeneratoruses the tool wgsim (https://github.com/lh3/wgsim) to simulate the sequence reads and concatenate them in a single fastq file. We set by default all the mutation and error rates to 0 but both can be modified in the source code.

### B Contribution matrix for Sec. 3.3

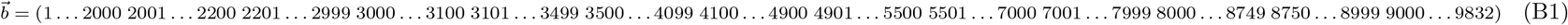

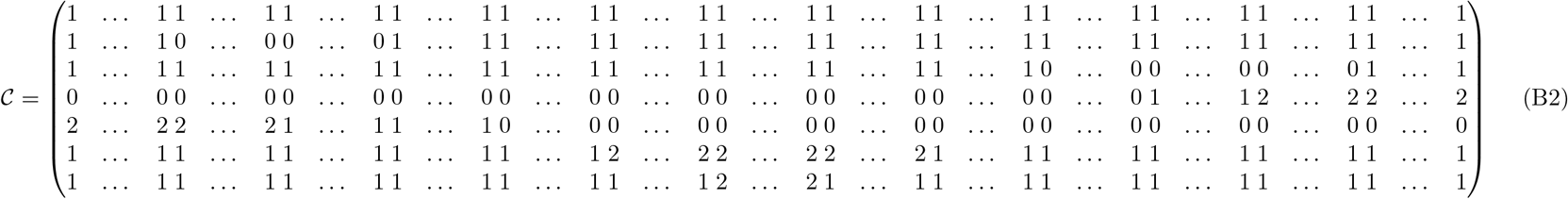

### C Pre-processing algorithm. DVGfinder output table curation

DVGfinder performs an intensive search on reads that do not match with the reference genome and classifies them into the most suitable category. This intensive search outcomes with a large table of candidate DVGs which needs to be curated. Here we present the pipeline we used to keep only those DVGs which events appear represented into the depth profile as a measurable change in the median of the plateaus adjacent to that DVG event. The pre-processing steps are the following:

i. Filtering Only those DVG candidates which number of reads conted by ViReMa-a outcomes a threshold, *R*_*th*_, are kept. Here we set this threshold relative to the maximum depth of the profile, 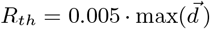. An example of the DVGfinder output table after this filtering step can be found at Table D1.
ii. Merging rows Rows which share the same DVG type and their positions differ below a certain threshold, Δ_*b*_, are merged. (here, Δ_*b*_ = 10). The resulting position will be the weighted mean (according to the number of ViReMa-a reads) of all merged rows. The associated number of reads will be the sum of the merged ones.
iii. Select event position with relevant change within plateaus’ medians Let’s call 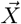 to the event position vector in which all DVG events (BP and RI) positions are stored. To avoid short plateau’s, positions in 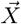 differing less than Δ_*b*_*/*2 are combined keeping the mean value. 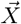 defines the limits of the plateaus. In this step we will remove all event positions which adjacent plateau’s median do not differ more than a certain threshold relative to the noise level.
  a. Obtain the noise level of the depth profile. The noise level computation follows the same procedure as in Sec. 3.2. Here, the noise level is computed as *n*_*l*_ = (|*u*_*b*_| + |*l*_*b*_|)*/*2, being *u*_*b*_ and *l*_*b*_ the upper and lower bounds of the relative depth deviation to the median.
  b. Compute the relative difference in consecutive plateau’s median values, 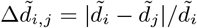.
  c. Keep those positions which plateau’s median depth at one side of the splitter and the other is greater than certain threshold relative to the noise level, 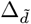. This is, keep positions in 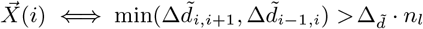.Here,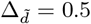.
iv. Filter DVGs without their BP or RI position present at the curated 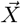 After having joined plateaus, it is possible that some DVGs do not have its start or ending position in 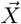. In this final step we will discard those DVGs as they not contribute to the depth profile. In this step, a threshold on the minimum event size, |BP−RI|, can be set.

The pre-procesing result reduces the number of DVGs of the sample from 8500 from the DVGfinder raw output to 34. The main reduction occurs during the filtering step. Fig. C1 show the depth profile with the event positions remaining after the pre-processing. Depending on the user’s motivation for the analysis, a different choice of parameters may be more appropriate. In case the goal is to have more power detecting DVGs, we will relax the parameters *R*_*th*_ and 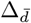. Finding the most restrictive parameters that still allow us to reconstruct this depth profile would be the most conservative option and most likely to give us DVGs actually present in the sample.

**Figure C1:**
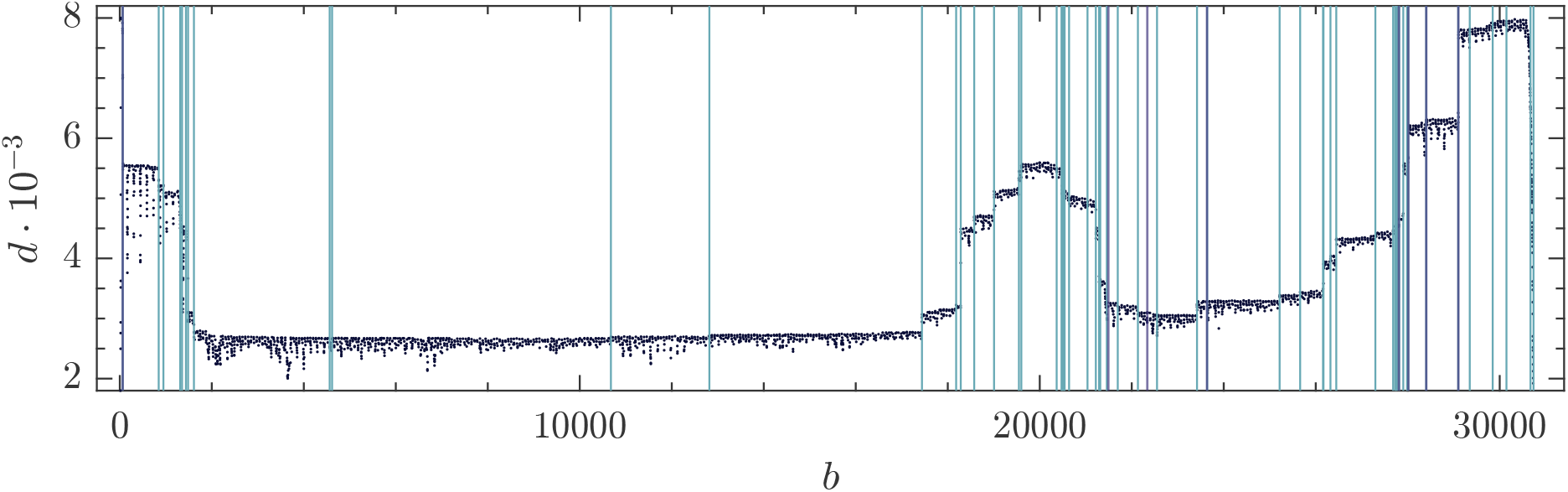
Depth profile with DVGs event’s positions indicated as blue lines. Purple lines indicate the localization of the known subgenomic positions for HCoV-OC43.

### D Case study: input data and results

**Table D1:**
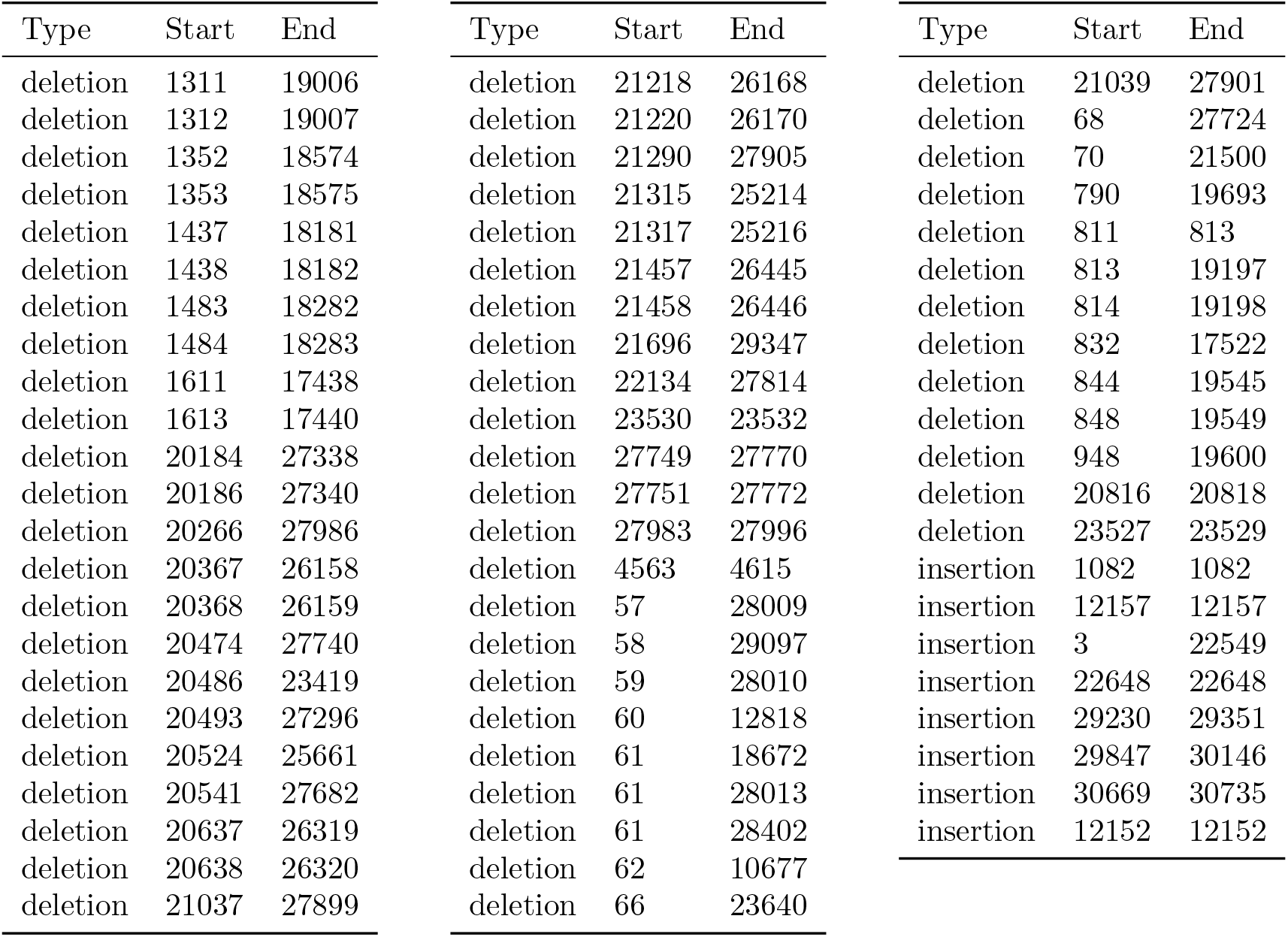
DVGfinder output tables using `metasearch` mode. DVG which appear less than *R*_*th*_ times, with *R*_*th*_ = 0.005 *·* 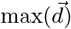, have been filtered.

| Type | Start | End | Type | Start | End | Type | Start | End |
| --- | --- | --- | --- | --- | --- | --- | --- | --- |
| deletion | 1311 | 19006 | deletion | 21218 | 26168 | deletion | 21039 | 27901 |
| deletion | 1312 | 19007 | deletion | 21220 | 26170 | deletion | 68 | 27724 |
| deletion | 1352 | 18574 | deletion | 21290 | 27905 | deletion | 70 | 21500 |
| deletion | 1353 | 18575 | deletion | 21315 | 25214 | deletion | 790 | 19693 |
| deletion | 1437 | 18181 | deletion | 21317 | 25216 | deletion | 811 | 813 |
| deletion | 1438 | 18182 | deletion | 21457 | 26445 | deletion | 813 | 19197 |
| deletion | 1483 | 18282 | deletion | 21458 | 26446 | deletion | 814 | 19198 |
| deletion | 1484 | 18283 | deletion | 21696 | 29347 | deletion | 832 | 17522 |
| deletion | 1611 | 17438 | deletion | 22134 | 27814 | deletion | 844 | 19545 |
| deletion | 1613 | 17440 | deletion | 23530 | 23532 | deletion | 848 | 19549 |
| deletion | 20184 | 27338 | deletion | 27749 | 27770 | deletion | 948 | 19600 |
| deletion | 20186 | 27340 | deletion | 27751 | 27772 | deletion | 20816 | 20818 |
| deletion | 20266 | 27986 | deletion | 27983 | 27996 | deletion | 23527 | 23529 |
| deletion | 20367 | 26158 | deletion | 4563 | 4615 | insertion | 1082 | 1082 |
| deletion | 20368 | 26159 | deletion | 57 | 28009 | insertion | 12157 | 12157 |
| deletion | 20474 | 27740 | deletion | 58 | 29097 | insertion | 3 | 22549 |
| deletion | 20486 | 23419 | deletion | 59 | 28010 | insertion | 22648 | 22648 |
| deletion | 20493 | 27296 | deletion | 60 | 12818 | insertion | 29230 | 29351 |
| deletion | 20524 | 25661 | deletion | 61 | 18672 | insertion | 29847 | 30146 |
| deletion | 20541 | 27682 | deletion | 61 | 28013 | insertion | 30669 | 30735 |
| deletion | 20637 | 26319 | deletion | 61 | 28402 | insertion | 12152 | 12152 |
| deletion | 20638 | 26320 | deletion | 62 | 10677 |  |  |  |
| deletion | 21037 | 27899 | deletion | 66 | 23640 |  |  |  |

**Table D2:** Result of the case study. *n*_*i*_ has been rounded to the nearest integer. Subgenomics appear colored in the table.

| Type | Start | End | $p_i(\%)$ | Reads | $n_i$ | Type | Start | End | $p_i(\%)$ | Reads | $n_i$ |
| --- | --- | --- | --- | --- | --- | --- | --- | --- | --- | --- | --- |
| deletion | 1312 | 19007 | 5.36 | 2916 | 421 | deletion | 21696 | 29349 | 0.98 | 46 | 77 |
| deletion | 1353 | 18575 | 2.61 | 1071 | 205 | deletion | 22134 | 27814 | 1.55 | 257 | 122 |
| deletion | 1438 | 18182 | 0.95 | 220 | 75 | deletion | 27750 | 27771 | 0.13 | 402 | 10 |
| deletion | 1484 | 18283 | 16.56 | 3066 | 1301 | deletion | 27985 | 27996 | 0.25 | 772 | 20 |
| deletion | 1612 | 17439 | 5.22 | 582 | 410 | deletion | 4563 | 4615 | 0.34 | 81 | 26 |
| deletion | 20368 | 26159 | 0.96 | 140 | 76 | deletion | 63 | 28010 | 8.42 | 1309 | 661 |
| deletion | 20474 | 27740 | 0.16 | 139 | 13 | deletion | 63 | 29098 | 18.91 | 5437 | 1486 |
| deletion | 20486 | 23419 | 2.87 | 920 | 225 | deletion | 63 | 12818 | 0.35 | 58 | 28 |
| deletion | 20493 | 27296 | 1.07 | 52 | 84 | deletion | 63 | 28402 | 1.23 | 201 | 97 |
| deletion | 20524 | 25661 | 0.60 | 100 | 45 | deletion | 63 | 10677 | 0.36 | 158 | 29 |
| deletion | 20541 | 27682 | 0.49 | 49 | 39 | deletion | 63 | 23640 | 0.25 | 71 | 20 |
| deletion | 20638 | 26320 | 1.12 | 900 | 88 | deletion | 63 | 27724 | 0.81 | 82 | 64 |
| deletion | 21038 | 27903 | 0.75 | 156 | 59 | deletion | 846 | 19547 | 3.96 | 1716 | 311 |
| deletion | 21219 | 26169 | 5.35 | 2852 | 421 | deletion | 948 | 19600 | 1.57 | 652 | 123 |
| deletion | 21290 | 27903 | 10.50 | 3603 | 825 | insertion | 1 | 22549 | 0.45 | 177 | 35 |
| deletion | 21316 | 25215 | 1.29 | 360 | 101 | insertion | 29847 | 30146 | 0.56 | 44 | 44 |
| deletion | 21458 | 26446 | 3.99 | 789 | 313 | insertion | 30669 | 30735 | $10^{-22}$ | 55 | 0 |

